# Single-cell genomics reveals region-specific developmental trajectories underlying neuronal diversity in the human hypothalamus

**DOI:** 10.1101/2021.07.20.453090

**Authors:** Brian R. Herb, Hannah J. Glover, Aparna Bhaduri, Carlo Colantuoni, Tracy L. Bale, Kimberly Siletti, Sten Linnarsson, Rebecca Hodge, Ed Lein, Arnold R. Kriegstein, Claudia A. Doege, Seth A. Ament

## Abstract

The development and diversity of neuronal subtypes in the human hypothalamus has been insufficiently characterized. We sequenced the transcriptomes of 40,927 cells from the prenatal human hypothalamus spanning from 6 to 25 gestational weeks and 25,424 mature neurons in regions of the adult human hypothalamus, revealing a temporal trajectory from proliferative stem cell populations to mature neurons and glia. Developing hypothalamic neurons followed branching trajectories leading to 170 transcriptionally distinct neuronal subtypes in ten hypothalamic nuclei in the adult. The uniqueness of hypothalamic neuronal lineages was examined developmentally by comparing excitatory lineages present in cortex and inhibitory lineages in ganglionic eminence from the same individuals, revealing both distinct and shared drivers of neuronal maturation across the human forebrain. Cross-species comparisons to the mouse hypothalamus identified human-specific *POMC* populations expressing unique combinations of transcription factors and neuropeptides. These results provide the first comprehensive transcriptomic view of human hypothalamus development at cellular resolution.

**One-Sentence Summary:** Using single-cell genomics, we reconstructed the developmental lineages by which precursor populations give rise to 170 distinct neuronal subtypes in the human hypothalamus.

## Main Text

The hypothalamus is a small but anatomically complex brain region that controls a large variety of evolutionally requisite physiological and homeostatic functions, including body temperature, circadian rhythms, sleep, stress responses, satiety, and hunger, and aspects of mood, social behavior, and memory(*1*–*3*). These functions are subdivided amongst specialized neuronal subtypes, which are organized into distinct anatomical nuclei(*4*–*11*). Environmental and genetic perturbations to hypothalamic development result in long-lasting changes in physiology and behavior(*12*–*20*) and are thought to contribute to risk for human diseases including obesity, anxiety, and depression(*21*, *22*). These clinical consequences motivate deeper investigation into the timing and regulation of hypothalamic development. However, much of what we know about hypothalamic development is derived from animal models(*2*–*12*, *23*, *24*). While many of these functions are thought to be evolutionarily conserved, the molecular identities of human hypothalamic cells and the timing and regulation of their development remain inadequately characterized(*25*, *26*). Here, we sought to address this deficiency through single-cell transcriptomics of the prenatal and adult human hypothalamus to define its transcriptional cell types and developmental trajectories.

## Results

### An atlas of neuronal and non-neuronal lineages in the developing human hypothalamus

We performed 10x Genomics single-cell RNA sequencing (scRNA-seq) of prenatal hypothalamus from 11 human fetuses (4 female, 7 male) at ~6 to 25 gestational weeks (GW); Carnegie Stages (CS) 13-15, CS22, GW16, GW18-20, GW22, GW25), yielding 40,927 high-quality single-cell transcriptomes (**Fig. 1A and tables S1**). These samples were collected in parallel with samples from several additional brain regions in the same fetuses, and scRNA-seq of cortical samples from these fetuses has been reported previously(*27*–*30*). In addition, we performed single-nucleus RNA sequencing (snRNA-seq) of neurons from the post-mortem hypothalamus of a neurotypical donor (male, 50 years of age) (**Fig. 1A and fig. S1**), yielding 22,365 high-quality single-nucleus transcriptomes.

**Figure 1.**
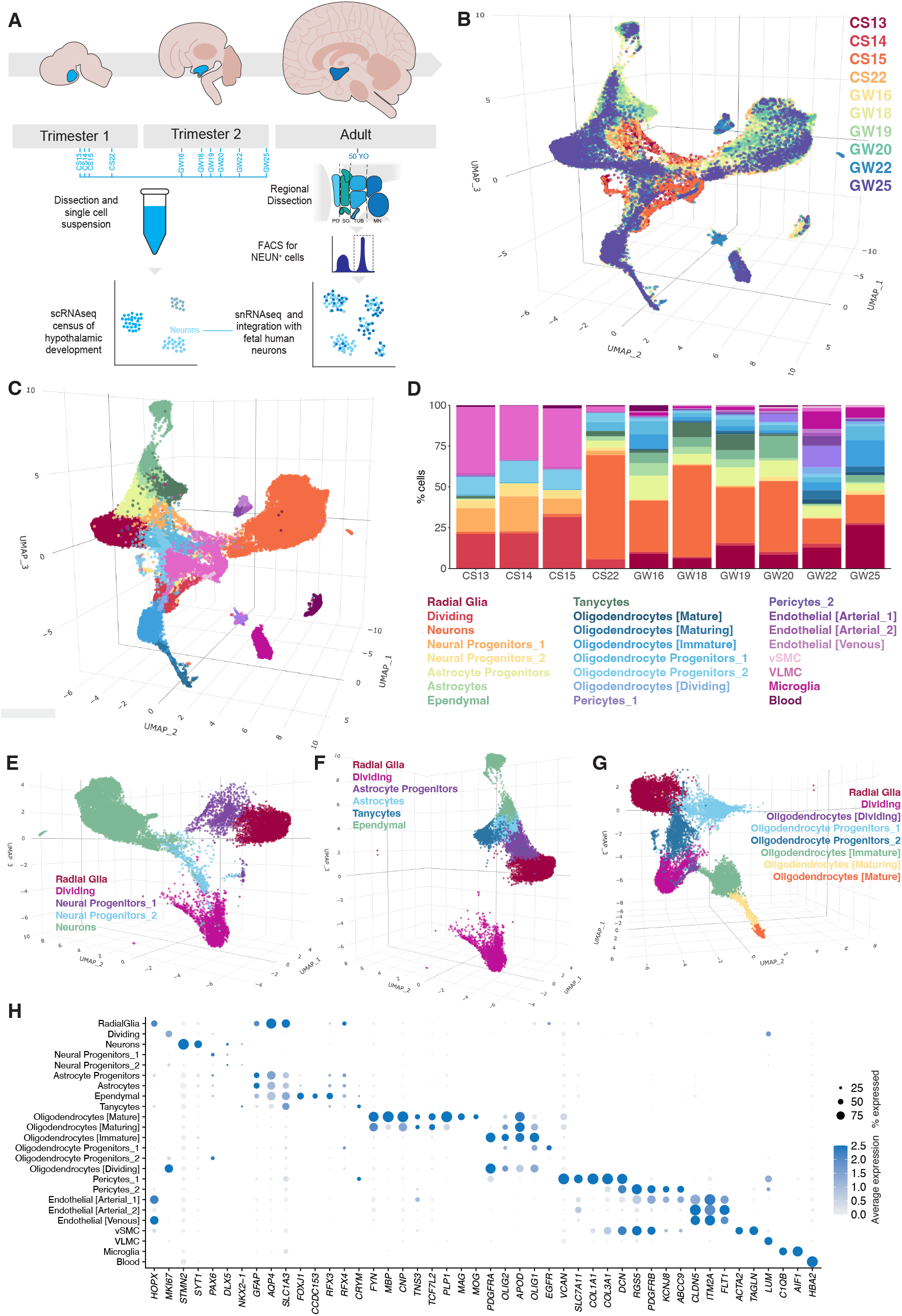
Neuronal and non-neuronal lineages across the developing human hypothalamus. A. Overview of sample collection, including single-cell RNAseq analysis of Trimester 1 and 2 prenatal hypothalamus, plus single-nucleus RNAseq from adult hypothalamic neurons dissected into four regions. B. <u>3D UMAP</u> showing integrated samples of the prenatal hypothalamic samples including Trimester 1 (CS13, CS14, CS15 and CS22) and Trimester 2 (GW15, GW18, GW19, GW20, GW22 and GW25). C. <u>3D UMAP</u> showing samples after clustering to show cell subpopulations. D. Stacked plot showing distribution of cell subpopulation across samples. E. 3D visualization of the <u>neuronal lineage</u>. F. 3D visualization of the <u>astro-ependymal lineage</u>. G. 3D visualization of the <u>oligodendrocyte lineage</u>. H. Dot Plot showing expression of marker genes.

We integrated the 40,927 prenatal hypothalamic cells (**Fig. 1B and fig. S2**) and annotated the resulting cell clusters using a curated, comprehensive list of literature-based marker genes (**tables S2 and S3**). We identified 24 broad cell classes, including radial glia, dividing cells, neuronal, oligodendrocyte, and astrocyte lineages, as well as other supporting cell types (**Fig. 1, C to H and table S4**). Cell type distributions shifted over developmental time (**Fig. 1, B to D**). The earliest timepoints – CS13, 14 and 15 – were composed largely of dividing progenitors (*MKI67^+^*) and vascular leptomeningeal cells (*LUM^+^*). Radial glia (*HOPX*^+^) cells were identified at GW16 and onward. Neuronal and oligodendrocyte progenitor populations were present from the earliest time point. Post-mitotic neurons (*STMN2^+^*) emerged as early as CS22 and were abundant starting at GW16. Oligodendrocyte populations (*PDGFRA^+^/OLIG2^+^/MKI67^-^*) were detected by GW16 and began to reach maturation (*MAG^+^/MOG^+^*) by GW22. Astrocytes (*GFAP^+^/AQP4^+^/HOPX^-^*), ependymocytes (*CCDC153*^+^), and tanycytes (*CRYM^+^*) were established by CS22.

Most neurons in the prenatal samples appeared immature, with weak expression of many canonical markers. To better annotate these populations and define their developmental trajectories, we integrated the 11,446 prenatal hypothalamic neurons (CS22 to GW25) with 22,365 adult hypothalamic neurons (**Fig. 2, A and B, and table S5**). Using Monocle3, we reconstructed a branching pseudotime trajectory, rooted at the earliest timepoint (CS22) (**Fig. 2, B and C**). We identified branch points where the lineages diverge, and ‘leaves’ where the lineages terminate and followed the trajectories of these branches and leaves to generate a lineage tree (**Fig. 2D**). The highest-order branch points distinguish GABAergic (*SLC32A1^+^*) and glutaminergic (*SLC17A6+*) neurons (**Fig. 2E**). At the next level, branches could primarily be assigned to specific hypothalamic nuclei (**Fig. 2, F and G and tables S6 and S7**). Ten hypothalamic nuclei were identified, including the Arcuate (ARC; *TBX3+*), Tuberomammillary Terminal (TM; *HDC*+), Paraventricular Nucleus of the Hypothalamus (PVH; *SIM1+*),Ventromedial Nucleus of the Hypothalamus (VMH; *FEZF1+*), Lateral Hypothalamus (LH; *LHX9*+), Suprachiasmatic Nucleus (SCN; *LHX8*+/*SIX6*+), Supramammillary Nucleus (SMN; *LMX1A+*), Mammillary Nucleus (MN; *FOXB1+*), Zona Incerta (ZI; *MEIS2+*) and Intrahypothalamic Diagonal (ID; *LHX6+*). The transcription factor signature for these nuclei could be detected by CS22 (first trimester), except for the SMN where robust expression could be detected in the second trimester (**Fig. 2, G to J and fig. S3**).

**Figure 2.**
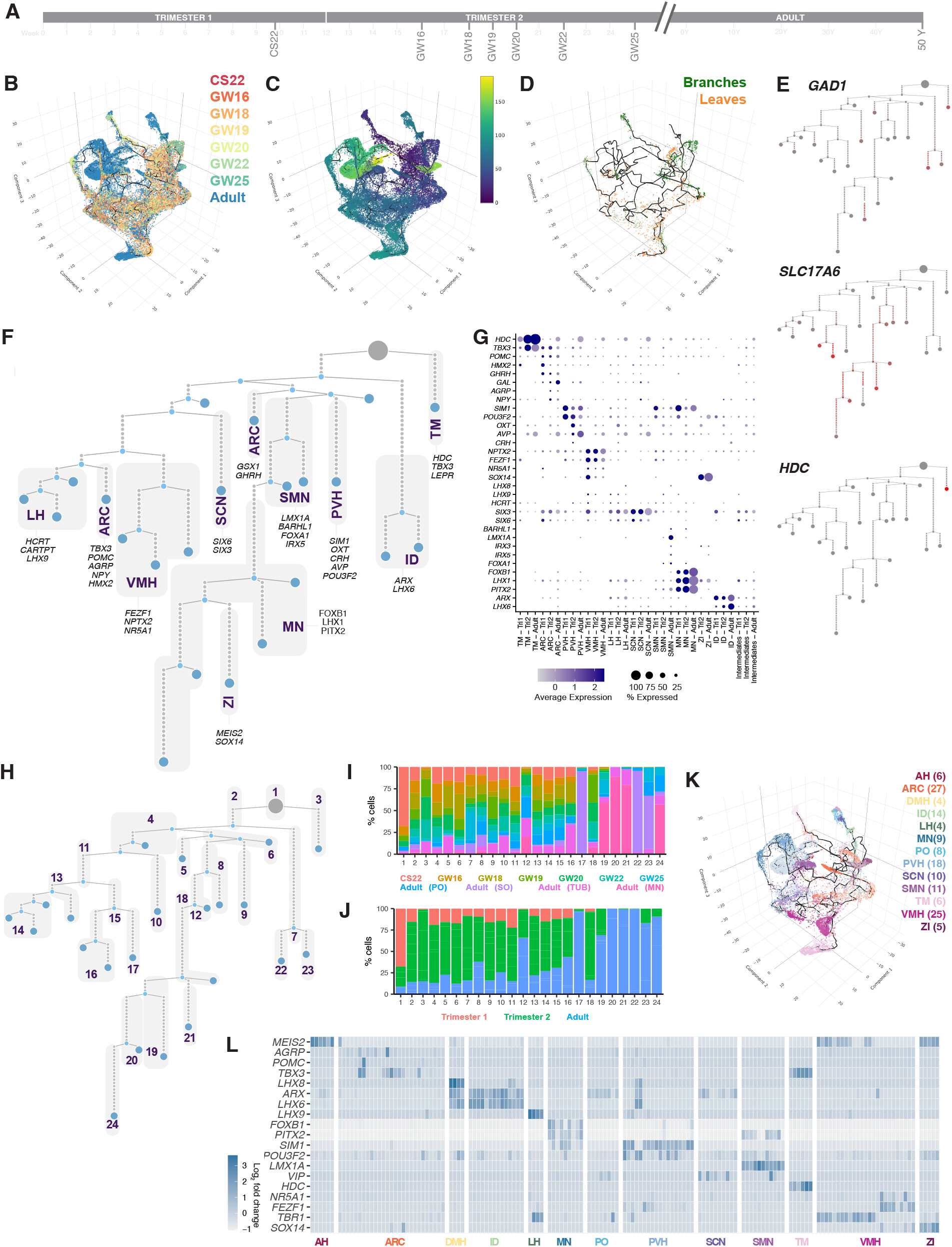
Neuronal trajectories across development and the adult human hypothalamus. A. scRNAseq data from embryonic postmitotic neurons (CS22-GW25) and neurons from the 50YO adult hypothalamus. B. <u>3D UMAP</u> showing overlay of samples by timepoint. C. Monocle3 was used to run <u>pseudotime</u> analysis of both prenatal and adult neurons starting with the node containing the highest percentage of cells from CS22. D. The <u>pseudotime trajectory</u> highlighting branch points (green) and the terminal leaves (orange) E. The pseudotime trajectory can be used to generate a 2D lineage tree. The lineage tree can be overlaid with gene expression (red). GABAergic (*GAD1*), glutaminergic (*SLC17A6*) and histaminergic (*HDC*) expression demarcates cell populations. F. Nuclei were annotated using the marker genes in fig. S4 and table S3. G. Dot Plot showing expression of marker genes and neuropeptides split between Trimester 1 and 2 and adult. H. The lineage tree was subdivided into 24 nodes to check sample distribution across tree. I. The 24 nodes were plotted to show the temporal distribution of nodes by time point. J. Temporal expression was also plotted to show distribution across trimester 1, 2 and adult samples. K. Hicat was used to generate <u>170 clusters</u> on the adult neurons. 147/170 clusters generated by Hicat could be assigned to nuclei. The number of individual nuclei are shown in parentheses, with nuclei visualized in shades within the Monocle3 lineage. L. Heatmap showing mean log2 fold change in expression of key marker genes for each individual subpopulation.

The expression of canonical hypothalamic neuropeptides demarcated the maturation of each nucleus, revealing variation across cell types. In the Arcuate nucleus, *POMC* and *GHRH* were expressed at low levels in the first semester and increased to robust levels by the second trimester. Other Arcuate neuropeptides such as *AGRP, NPY* (co-expressed in AGRP neurons), and *CARTPT* (co-expressed in POMC neurons) were expressed only starting in the second trimester. *AVP*, a canonical neuropeptide of the PVH could be detected by CS22, while other PVH neuropeptides such as *OXT* and *CRH* were not expressed until the second trimester (**Figure 2G**).

We applied a hierarchical iterative clustering approach (scrattch.hicat package) to obtain a more detailed atlas of neuronal subtypes. This revealed 170 transcriptionally distinct cell clusters in adult neurons. 147 of which could be mapped to a specific nucleus (**Fig. 2, K and L)**, including all ten nuclei represented in our lineage tree and three additional nuclei that were more sparsely represented in the data **(fig. S4 and tables S3 and S8**). 74 clusters were glutaminergic, 67 were GABAergic and 6 were histaminergic (**table S9**). Most clusters could be assigned to established neuronal subtypes by the expression of neuropeptides and other canonical markers. For instance, we identified discrete neuronal populations expressing POMC, AGRP, OXT, GHRH, and AVP, each of which was restricted to just 1-2 clusters. Our data also support diverse subpopulations expressing certain neuropeptides, including CRH, and TRH, and SST, each of which was expressed in ten or more clusters. In summary, our data define the development and diversity of neuronal subtypes in the human hypothalamus in unprecedented detail.

### Cross-species comparison of neuronal subtypes in humans and mice

Major classes of hypothalamic neurons are shared in humans and mice, but species differences in more refined neuronal subtypes are poorly understood. We integrated our sc/snRNA-seq data from human prenatal and adult hypothalamic neurons with existing scRNA-seq from developing(*11*) and adult mouse hypothalamus(*4*–*6*, *8*, *9*, *11*) to create a comprehensive atlas of hypothalamic neurons (*n* = 175,754, **Fig. 3A and table S10**). As expected, human and mouse neurons co-clustered within major groups corresponding to immature (prenatal) neurons, GABAergic neurons, glutamatergic neurons, and histaminergic neurons (**Fig. 3, B and C**). Louvain clustering revealed 28 neuronal sub-classes, most of which could be annotated to established subtypes and nuclei (**Fig. 3, D and E and fig. S5** and **table S11**). Most sub-classes contained both human and mouse cells, consistent with the evolutionary conservation of most hypothalamic neuronal sub-classes. While certain clusters of adult neurons appeared more abundant in one species than the other, we believe this most likely results from sampling differences in the human vs. mouse datasets rather than true species differences at this sub-class level. Thus, our analysis supports dozens of evolutionary conserved neuronal subclasses and describes their marker genes.

**Figure 3.**
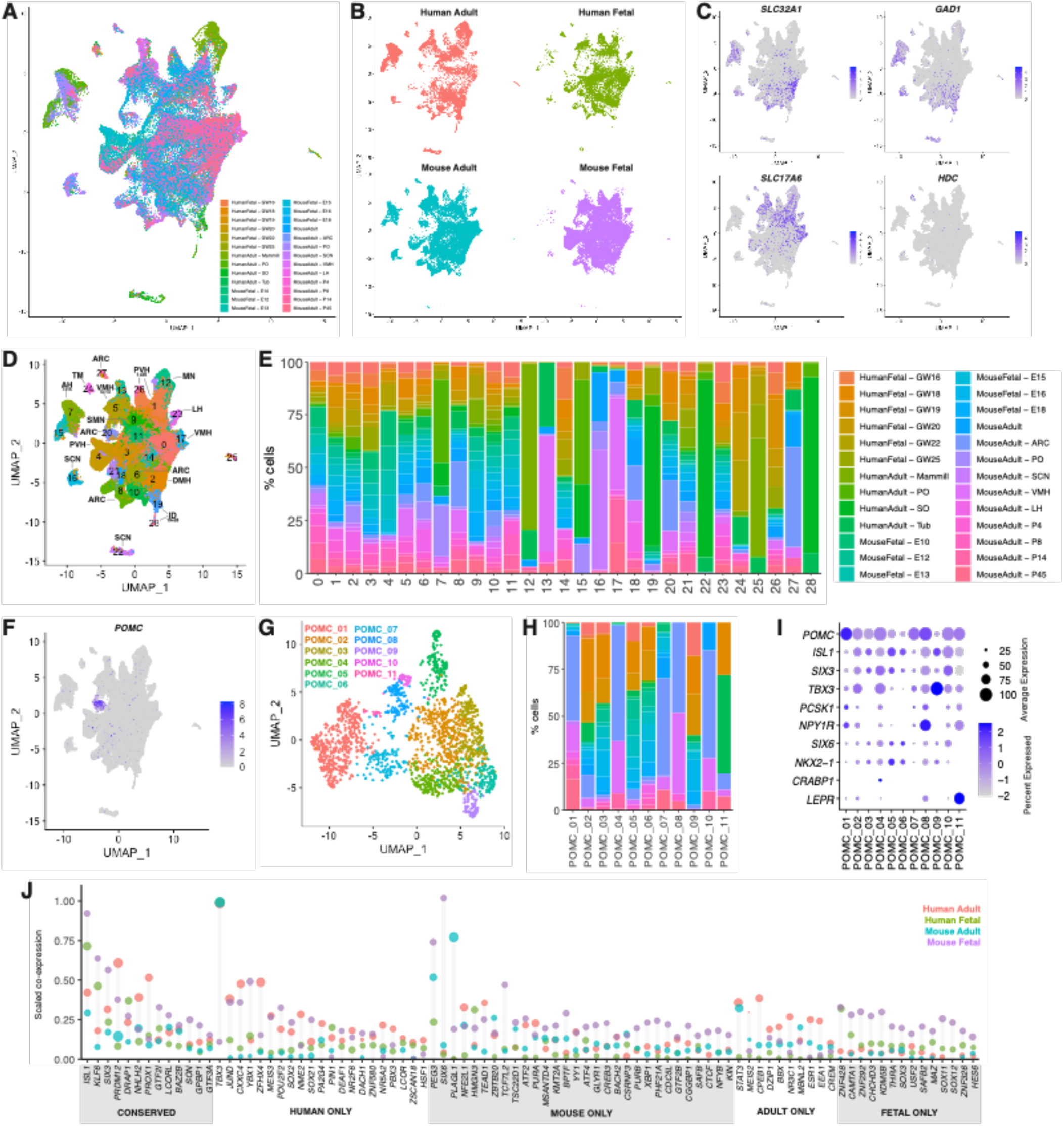
Cross-species comparison of mouse and human hypothalamic neurons. A. UMAP of single-cell RNAseq data from the human and mouse hypothalamus neurons, including prenatal and adult samples. Mammill, mammillary dissection; PO, preoptic dissection; SO, Supraoptic dissection; Tub, Tuberal dissection, ARC, Arcuate; SCN, Suprachiasmatic Nucleus; VMH, Ventromedial Nucleus of the Hypothalamus; LH, Lateral Hypothalamus. B. Split UMAP showing cell distribution across human/mouse and prenatal/adult samples. C. FeaturePlots showing localization of GABAergic (*SLC32A1* and *GAD1*), Glutaminergic (*SLC17A6*) and histaminergic (*HDC*) neurons. D. Louvain clustering generates 28 cell clusters. Anatomical nuclei annotations are indicated for clusters based on sample dissection and gene expression profile. E. Stacked plot showing the percentage of samples contributing to each cluster. Percentage is normalized to the number of cells in the input, with legend shown on right. F. FeaturePlot showing localization of *POMC* neurons. G. Subclustering of POMC neurons reveals 11 unique POMC^+^ clusters. H. Stacked plot showing the percentage of input samples contributing to each POMC cluster. Percentage is normalized to the number of cells in the input, with legend shown Fig. 3E above. I. DotPlot showing expression of genes localized within POMC subclusters. J. Transcription factor co-expression analysis across humanImouse and fetalIadult samples. Transcription factors coexpressing with POMC cells with a co-expression value three standard deviations above the mean are highlighted for each category. Dot size indicates relative the expression level for the gene of interest.

To explore species differences in more refined sub-populations, we focused on POMC neurons, which were well represented in both the human and mouse datasets (**Fig. 3F**). Subclustering of POMC neurons revealed 11 sub-clusters (**Fig. 3G and tables S12 and S13**). Overall, cells from mouse and human samples were well mixed (**Fig. 3I**). However, cluster 11 (*POMC/PCSK1/LEPR/NPY1R/TBX3+*) was over-represented for human cells (79% human), while clusters 1, 4, 7, 8 and 10 were over-represented for mouse cells (95%, 98%, 82%, 100% and 100% respectively) (**Fig. 3, H to J**). We used co-expression patterns of transcription factors and neuropeptides within POMC neurons to gain insight into both conserved and species-specific regulatory networks (**tables S14 to S18**). This analysis predicted a core set of TFs as master regulators shared by mouse and human POMC neurons, including *ISL1, KLF6, SIX3, PRDM12, NHLH2* and *PROX1*. However, human POMC neurons also utilize *TBX3, JUND, CXXC4, YBX1, ZFHX4*, *MEIS3*, and *POU2F2*, whereas mouse POMC neurons utilize *PEG3*, *SIX3*, *PLAGL1* and *NFE2L1* (**Fig. 3K**). Thus, our data provide preliminary evidence for human-specific POMC neuron subtypes and their regulatory networks, which may merit further investigation.

### Unique and shared features of hypothalamic vs. cortical germinal zones in the human forebrain

Little is known about the genes and trajectories distinguishing neurogenesis in the hypothalamus versus other forebrain regions. To address this, we obtained matched samples from the cortex, ganglionic eminence (the source of telencephalic inhibitory neurons), and hypothalamus from the same individuals at GW18, GW19 and GW20. Co-embedding these samples in a shared low-dimensional space enabled direct comparisons among the progenitor populations and excitatory and inhibitory neuron lineages (n=95,107, **Fig. 4 A and B and fig. S6 A and B**). Both neuronal and non-neuronal lineages emanate from a large cluster of radial glia that were abundantly represented in all three stem cell niches. Radial glia give rise to multiple neuronal and numerous non-neuronal progenitor populations. Of these, among the non-neuronal populations, we identified tanycytes and ependymal cells that were specific to the hypothalamus, as well as astrocytes and oligodendrocytes that were abundant in both the hypothalamus and cortex. A large population of dividing progenitor cells formed a distinct barrel shape in 3D UMAP space indicative of actively cycling cells (cell cycle stage, **fig. S6C**, seurat clusters 6, 9, 10, **fig. S6 D**). These dividing progenitors, along with the radial glia, feed into a neuronal lineage that split between excitatory and inhibitory lineages and further subdivided into major neuron sub-classes. While cortex and GE samples split mid-way along the developmental lineage between progenitors and mature neuron populations in a largely expected way, we took a keen interest in similarities and differences among these lineages and how progenitors from the hypothalamus fit into this framework.

**Figure 4.**
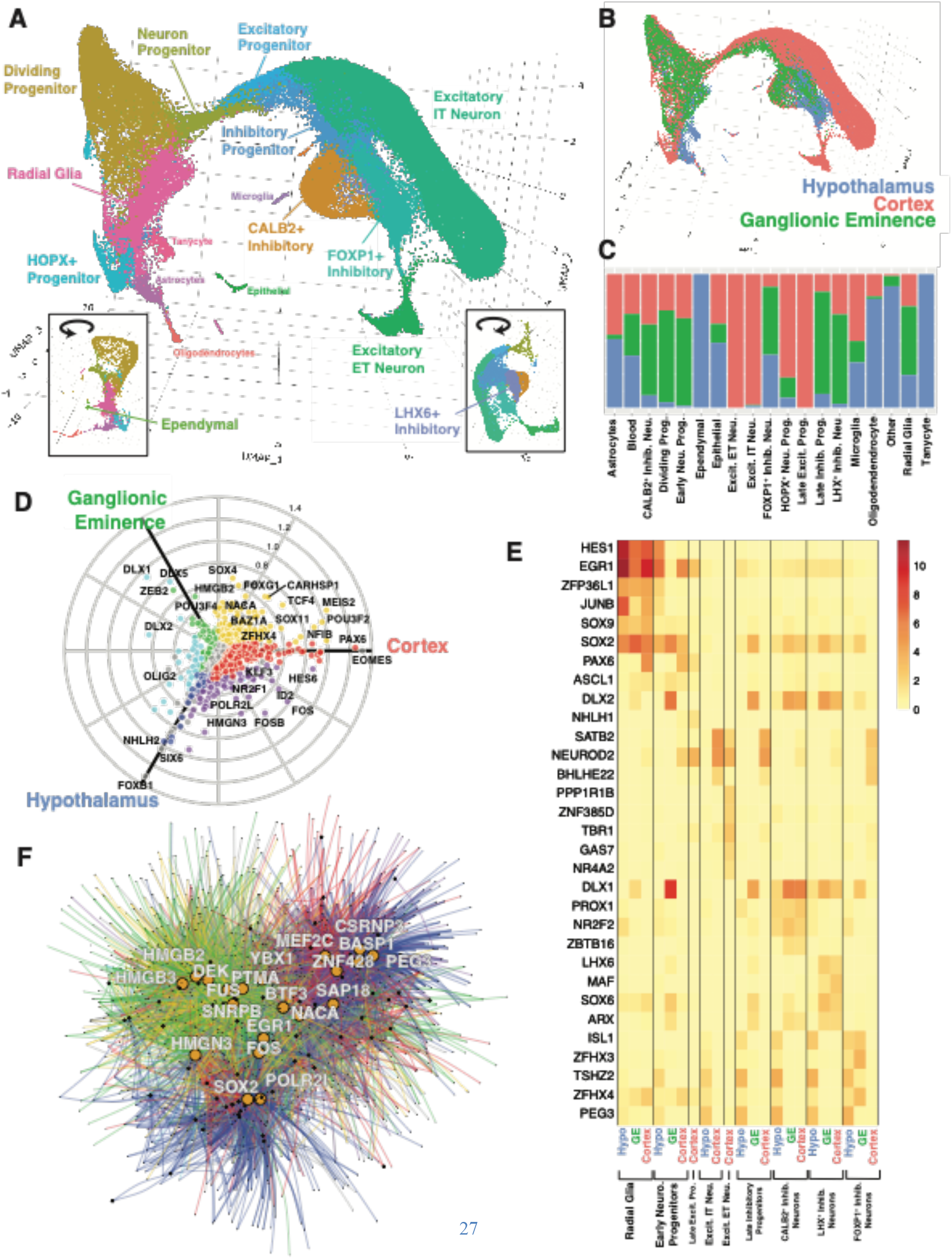
Developmental principles across distinct human brain regions. A. scRNAseq data from hypothalamus, cortex and ganglionic eminence for three matched human embryonic samples at GW18, GW19 and GW20. Cell types presented in <u>3D UMAP</u>, inset panels are rotations of UMAP showing additional clusters. B. <u>3D UMAP</u> showing overlay of samples by brain region. C. Distribution of cells within cell-type cluster by brain region. D. Polar plot of TF expression across three brain regions for dividing progenitor cells. Points off-axis reflect shared expression between regions. Grey points indicate no significant difference between regions. E. Heatmap depicting genes with similar or divergent expression patterns across neuronal lineages. Colors represent scaled expression of cells only within a given cell-type and brain region, as described in x-axis legend. F. Subsets of gene regulatory networks for hypothalamus, cortex and ganglionic eminence were created based on genes expressed in neural progenitor cells, radial glia and dividing progenitors. Resulting overlapping networks show edges from hypothalamus network (blue) are roughly evenly split and adjacent to the ganglionic eminence network (green) and cortex network (red). Shared edges follow the color scheme introduced in Fig. 4D. Top 20 TFs with the greatest number of connections are highlighted in white.

The co-embedded neurogenic map retains well-known features of cortical neurogenesis. Excitatory neurons of the cortex branched off from inhibitory neurons about halfway along the bridge from early neuron progenitors into mature neuron populations. This lineage led directly into intratelencephalic (IT) excitatory neuron populations and reflected the temporal ordering with which deep layer vs. superficial layer IT neurons are born: newborn neurons at the beginning of the lineage proximal to progenitor cells primarily expressed markers for later-born superficial layer excitatory neurons such as *CUX2*, while more mature neurons at the lineage terminus primarily expressed markers of earlier-born deep layer 5/6 neurons such as *RORB*. A distinct branch in the trajectory expressed markers of extratelencephalic (ET) excitatory neurons. Separately, inhibitory neuron lineages led into three distinct inhibitory neuron clusters, which expressed markers of medial ganglionic eminence-derived (*LHX6+*), caudal ganglionic eminence-derived (*CALB2^+^*), and lateral ganglionic eminence-derived (*FOXP1^+^*) subpopulations. The hypothalamic germinal zone is distinct from germinal zones of the cortex and GE in that it gives rise to both excitatory and inhibitory neurons, and these locally born neurons represent most of the mature neurons within the hypothalamus. Most of the hypothalamic cells were distributed among excitatory and inhibitory lineages that also included cortical and GE-derived cells. This suggests that at a high level of organization the development of excitatory vs. inhibitory neurons in the hypothalamus involves shared regulatory programs with the development of excitatory vs. inhibitory lineages in the cortex and GE.

The high-level similarity of the developmental trajectories in the cortex, GE, and hypothalamus enabled direct comparisons of the lineages to reveal finer-scale distinctions. Region-specific gene expression differences were already apparent at early stages of development. In radial glia and actively dividing progenitor populations (clusters 4, 6, 9, 10), we found region-specific expression for developmental transcription factors such as *PAX6* and *POU3F2* in the cortex, *DLX1* and *DLX5* in the GE, and *FOXB1* and *SIX6* in hypothalamus (**Fig. 4D and tables S19 and S20**). We also identified shared regulators, including *HES1* and *EGR1* (**Fig. 4E**). At a later stage of development, in neural progenitors (cluster 11), we found hypothalamus-specific expression for the transcription factor *TSHZ2*, which remained elevated in mature neuron populations. GE-specific transcripts in neural progenitors included *COL1A2*, a collagen, along with elevated expression of the NOTCH signaling ligand *DLL1* and the cell-cell signaling mediator *LGALS1*. We confirmed cortex-specific expression for canonical progenitor marker genes such as *EOMES* and *NEUROD2*. Many other genes showed complex spatiotemporal patterns in neural progenitors. For instance, neural progenitors of the hypothalamus and cortex had high expression (compared to GE) for cytoskeletal proteins such as *MYO10, CALD1*, and *VIM*, whereas neural progenitors in the hypothalamus and GE (but not cortex) had shared expression of *RBP1*, involved in retinoic acid signaling(*31*). Other genes had different temporal patterns or different cell type specificity. For example, *PPP1R17*, a phosphatase regulatory subunit, was identified as a neuroprogenitor marker within the cortex(*30*). However, in the hypothalamus, *PPP1R17* is instead expressed most highly in radial glia. These results provide insight into the unique regulatory programs that give rise to specialized neuronal populations.

We applied a network-biology approach to gain further insight into the shared vs. unique drivers of neuronal development across lineages. Gene co-expression network modeling identified 28 gene co-expression modules, many of which were expressed in region- and stage-specific patterns across the development of excitatory and inhibitory neuronal lineages (**fig. S6E**). In addition, we reconstructed a gene regulatory network (GRN) model(*32*) to predict the target genes for 1317 TFs in each germinal zone and the key regulator TFs for each gene coexpression module. We found both region-specific and shared TF-gene associations, where the extent of overlap across models provides a measure for the re-wiring of these networks across brain regions. We illustrate this concept for the 20 most highly-connected ‘hub’ transcription factors (**Fig. 4F**). For these TFs, we detect many TF-gene edges that are reproducible within different samples from the same brain region but not across brain regions, suggesting substantial re-wiring. Interestingly, network edges detected in the hypothalamus appear to be split between cortical and GE-derived groups, suggesting an early lineage difference not distinguished in the UMAP in **Fig. 4A**. Gene network modeling also provided insights into the regulation of specific neuronal lineages. The gene co-expression module M73 (**fig. S6E**) was expressed across the entire excitatory neuron lineage from dividing progenitor cells to mature neurons, enabling us to compare the regulators of excitatory neuron development across regions. Several TFs were identified as key regulators of excitatory neuron development in both hypothalamus and cortex, including *NEUROD6, ANK2, MYT1L*, and *CSRNP3*. Strikingly, we also identified regionspecific regulators of excitatory neurons, including hypothalamus-specific (e.g., *CSRNP3, ARID4A, PEG3, BASP1*) and cortex-specific TFs (*MEF2C, ZBTB18, NEUROD2, SATB2*). Similarly, module M70 was expressed across the entire inhibitory lineage, and we used this module to compare the regulators of inhibitory lineages in hypothalamus and GE (**fig. S6E**). We identified *SP9, DLX5, DLX2, SOX11*, and *SOX4* as shared regulators of inhibitory neuron development across regions, in addition to hypothalamus-specific (*SCAND1*, *ZNF428*, *YBX1*, and *PTMA*) and GE-specific TFs (*CITED2, TCF4, LHX6*, and *ARX*). This same pattern of shared and region-specific regulators could be identified in several additional modules, including M42, which is activated at later stages of excitatory neuron development (**fig. S6E**). These results suggest that there are shared “core” regulatory programs that govern the development of excitatory and inhibitory neurons in multiple brain regions, which may be refined by regionspecific regulatory programs that give rise to the unique properties of each region’s excitatory and inhibitory populations.

## Discussion

Here, we have described the development and diversity of neuronal and non-neuronal cell types of the human hypothalamus across prenatal stages to adulthood at single-cell resolution. The window into prenatal development starting at ~6 GW and including 25 GW is the largest to date, and our data represent the first single-nucleus transcriptomic atlas for the adult human hypothalamus. Our comparisons of hypothalamic cells in humans vs. mice expands knowledge about both shared and human-specific aspects of hypothalamic development. Comparisons among matched prenatal brain samples provided a unique opportunity to evaluate specializations of neurodevelopmental trajectories in the hypothalamus vs. other forebrain regions.

Our analysis revealed a sequential specification of hypothalamic neurons. First, developing neurons attain a unique transcriptional identity corresponding to the nucleus in which they reside. Later, specific neuronal subtypes differentiate. This process was well underway by Carnegie Stage 22, just ~10 weeks of gestational age, at which time we were able to discern most nuclei by the expression of established markers. We confidently identified ten hypothalamic nuclei in our human prenatal data, and human cells from three more nuclei could be identified after including complementary data from human adult. Notably, certain nuclei appeared to mature at different rates. Distinct developmental dynamics among hypothalamic nuclei are consistent with previous reports using immunohistochemistry(*33*) as well as a recently published single-cell transcriptomics study of the human prenatal hypothalamus covering GW7 - 20(*34*).

Differentiation of neuronal subtypes occurred primarily in the second trimester samples, with evidence that the maturation of many hypothalamic neurons continues into the third trimester and beyond. Our results confirm previous observations that *OXT* is detectable by GW14(*35*–*37*) and *CRH* and *AVP* as early as GW12(*38*), although we observe *AVP* as early as GW10. However, even late in the second trimester, many neuronal subtypes were identifiable primarily by the expression of key transcription factors, while the expression of some neuropeptides remained sparse, such as *KISS1*. Such sparse expression of neuropeptides in the second trimester might be interpreted as still ongoing differentiation, diversification, and maturation of the various peptidergic subtypes.

We detected 170 transcriptionally distinct neuronal populations in the adult hypothalamus, 147 of which could be mapped to specific nuclei. These numbers are within the range of detected neuronal cell types by scRNA-seq at postnatal timepoints in mice(*4*–*11*), and more than three times the number of distinct populations described previously for the human hypothalamus(*34*). The increased resolution of our dataset can be attributed to adult timepoints with more mature neurons. However, future large-scale studies will be needed to better define some of the rare subtypes.

As noted above, variation in the prenatal environment can have lasting consequences on many hypothalamic functions(*12*, *13*, *22*), but it is not well understood why adverse prenatal environments lead to worse or different outcomes in some individuals and not others. Precisely delineating the timing at which hypothalamic nuclei mature could provide insight into these exposures and outcomes, including the possibility that differences in the developmental timing among hypothalamic nuclei could produce distinct sensitive periods within the developmental trajectory and different disease risks.

Mouse models have been used extensively to study the relationships between the prenatal environment, the development of the hypothalamus, and the emergence of behavioral and physiological variation in hypothalamic functions. Thus, a critical question is whether these developmental processes are strongly conserved in mice vs. humans. Overall, we found that patterns of cell type-specific gene expression were quite similar in mouse vs. human. Known markers from mice enabled us to assign nucleus and cell type identities for all nuclei and many neuronal subtypes(*11*, *39*–*46*). However, numerous human-specific gene expression patterns were identified, including human-specific utilization of certain transcription factors, which could provide a substrate for subtle changes in regulation and function. Therefore, as has been demonstrated in other brain regions, many cell type-specific gene expression programs for neuronal subtypes are conserved from mice to humans. For example, among the conserved genes are the transcription factors *ISL1* and *NHLH2* which have been implicated in the regulation of body weight via hypothalamic circuitry in mice and humans(*41*, *47*–*49*). Nonetheless, results in mice should be interpreted cautiously when specific genes show divergent patterns across species.

Our comparison of forebrain neurogenic niches revealed shared lineages leading to excitatory and inhibitory neurons in the hypothalamus, cortex, and GE. However, gene expression differences were detectable between regions starting at early stages of development in radial glia. Gene regulatory network models suggested that the differentiation of excitatory vs. inhibitory neurons occurs early in hypothalamic development and involves ‘core’ regulatory programs that are shared, respectively, by excitatory and inhibitory neuronal lineages in the cortex and GE. We present evidence that these core programs are refined by region-specific TFs that may give rise to the unique properties of the excitatory and inhibitory programs in each brain region. While some of these predicted regulators have been described previously, our models make many novel predictions for the roles of key regulator TFs, and these should be tested experimentally. As some of these networks are predicted to be human-specific, testing their functions will likely require the further development of human cell culture systems that enable direct comparisons across forebrain stem cell niches.

Several limitations should be noted, setting the stage for future work. As with most human studies, information on the third trimester is lacking. Additional scRNA-seq data from late gestation and early postnatal time points could improve the resolution to characterize the maturation of nuclei and cell types. In addition, there is a need for more spatially-resolved human transcriptomic data to more fully annotate neuronal subtypes to the nuclei in which they reside. Finally, there is limited information about the prenatal environment experienced by the fetuses that we studied. An exciting future direction will be to expand our analysis to human and model organism samples with known variation in the prenatal environment, enabling a more direct evaluation of sensitive periods and their developmental consequences. It is our expectation that continuing to characterize human-specific patterns of gene expression in the hypothalamus will enable researchers to more precisely evaluate genes and hypothalamic cell types in the pathophysiology of human diseases.

## Acknowledgments

We thank Owen White and the Neuroscience Multi-Omic Archive team and Ronna Hertzano and the gEAR / NeMO Analytics team, respectively, for their assistance in building and hosting the data resources and web browsers that accompany this study. Some primary human tissue was obtained from the Human Developmental Biology Resource (HDBR), with special thanks to S. Lisgo and M. Crosier.

## Author contributions

Conceptualization: BRH, HJG, TLB, CAD, SAA

Methodology: BRH, SAA, CC

Investigation: BRH, HJG

Resources: AB, KS, SL, RH, EL, ARK

Supervision: CAD, SAA

Writing – original draft: BRH, HJG, CAD, SAA

Writing – review & editing: all authors

## Data and materials availability

The data analyzed in this study were produced through the Brain Initiative Cell Census

Network (BICCN:RRID:SCR_015820) and deposited in the NEMO Archive (RRID:SCR_002001) under identifier nemo:dat-917e9vs accessible at https://assets.nemoarchive.org/dat-917e9vs.

Data can be explored on NeMO Analytics: https://nemoanalytics.org/p?l=a856c14e&g=gad2

## Code availability

https://github.com/brianherb/HumanHypothalamusDev

## Supplementary Materials

Materials and Methods

Figs. S1 to S6

Tables S1 to S24

## Materials and Methods

### Prenatal sample collection and processing

Acquisition of all primary human tissue samples was approved by the UCSF Human Gamete, Embryo and Stem Cell Research Committee (approval nos. 10-03379 and 10-05113). All experiments were performed in accordance with protocol guidelines. Informed consent was obtained before sample collection and use for this study. First and second trimester human hypothalamus tissue was collected from elective pregnancy termination specimens from San Francisco General Hospital and the Human Developmental Biology Resource (HDBR). Cortical and ganglionic eminence tissue was collected in parallel from the same specimens, as previously described(*29*, *30*). Hypothalamic tissue samples were dissociated using Papain (Worthington) containing DNase. Samples were grossly chopped and then placed in 1mL of Papain and incubated at 37C for 15 min. Samples were inverted three times and continued incubating for another 15 min. Next, samples were triturated by manually pipetting with a glass pasteur pipette approximately ten times. Dissociated cells were spun down at 300g for 5min and Papain removed.

### Adult sample collection and processing

Adult hypothalamus tissue was obtained by the Allen Institute from a 50 year old Male donor (subject ID H18.30.002, Right hemisphere, Cause of death = Cardiovascular, PMI = 10hr, Tissue RIN score = 8.2 +/− 0.4) with no history of neuropsychiatric or neurological disorders and negative for infectious disease. Tissue collection was performed in accordance with the provisions of the United States Uniform Anatomical Gift Act of 2006 described in the California Health and Safety Code section 7150 (effective 1/1/2008) and other applicable state and federal laws and regulations. The Western Institutional Review Board reviewed tissue collection procedures and determined that they did not constitute human subjects research requiring institutional review board (IRB) review. The de-identified postmortem brain sample was obtained after receiving permission from the decedent’s legal next-of-kin and prepared as described previously (full methods described in Hodge 2019(*50*) (dx.doi.org/10.17504/protocols.io.bf4ajqse) and Bakken 2021(*51*) describes subject ID H18.30.002 sample). Briefly, coronal brain slabs of 1cm thickness were frozen in dry-ice cooled isopentane, and transferred to vacuum-sealed bags for storage at −80°C until the time of further use. To isolate the brain regions of interest, tissue slabs were briefly transferred to −20°C and the region of interest was removed and subdivided into smaller blocks on a custom temperature controlled cold table. Tissue blocks were stored at −80°C in vacuum-sealed bags until later use. **fig. S1** details the four broad regions – preoptic, supraoptic, tuberal and mammillary, identified by gross anatomical landmarks for the hypothalamus dissections. Nucleus isolation for 10x Chromium Single Cell 3’ RNA sequencing V3 was conducted as described (dx.doi.org/10.17504/protocols.io.y6rfzd6). Gating on DAPI and NeuN fluorescence intensity was carried out as described previously(*50*). NeuN+ and NeuN-nuclei were sorted into separate tubes and were pooled at a defined ratio after sorting. Sorted samples were centrifuged, frozen in a solution of 1X PBS, 1% BSA, 10% DMSO, and 0.5% RNAsin Plus RNase inhibitor (Promega, N2611), and stored at −80°C until the time of shipment on dry ice from the Allen Institute to the Karolinska Institute for 10x chip loading.

### Sequencing

For prenatal samples, single-cell capture was performed following the 10X v2 Chromium manufacturer’s instructions. Each sample was its own batch. For each batch, 10,000 cells were targeted for capture and 12 cycles of amplification for each of the complementary DNA and library amplifications were performed. Libraries were sequenced according to the manufacturer’s instructions on the Illumnia NovaSeq 6000 S2 flow cell (RRID:SCR_016387). For adult samples, immediately before loading on the 10x Chromium instrument, frozen nuclei were thawed at 37°C, washed, and quantified for loading as described (dx.doi.org/10.17504/protocols.io.nx3dfqn). In brief, suspensions were thawed in a 37°C water bath, spun down briefly, and pipetted several times to mix. Nuclei were then processed according to the 10X Genomics protocol, targeting 5000 cells. We aimed to sequence two replicates per sample. Nearly all samples were processed with the 10X Genomics V3 kit. Samples were first sequenced to a shallow depth (approximately 1000 reads/cell) on the Illumina NextSeq platform to validate sample concentrations. Samples were then sequenced to approximately 100,000 readsIcell on the Illumina NovaSeq platform. After sequencing, saturation was calculated for each sample using the Preseq package (https://github.com/smithlabcode/preseq). Any samples that were not saturated to 60% were sequenced more deeply using Preseq predictions.

### Quality filtering and integration

All fetal samples were quality filtered to include cells with the number of detected genes between 200 & 4000, a total UMI between 1,000 and 15,000 and a maximum 10 percent of reads mapping to mitochondrial genes. Adult neurons were filtered to include cells with at least 500 reads, and total UMI above 1000. For published datasets, cells passing quality filters defined in publication were used in this study. Doublets were detected using scDblFinder (default parameters, clust.method=‘overcluster’) and discarded from the analysis. Unless otherwise noted, the *Seurat* package v4.0.5(*52*) was used for normalization, sample integration, clustering, differential gene expression analysis and plotting. Prior to integration, samples were normalized using SCT transform, regressing on percent mitochondrial content.

### Cell type assignments

Integrated prenatal samples (n cells = 40,927) were subject to dimension reduction with PCA and were visualized in the 3D UMAP space. Cells were clustered using Louvain clustering and identified using established markers from published studies(*2*, *4*–*11*, *23*, *53*–*57*). Aberrant cells were manually reassigned based on UMAP coordinates to best reflect gene expression in feature plots. Extrahypothalamic cells such as *FOXG1+* cells (Telencephalon) and *NEUROD6+* cells (forebrain) were removed prior downstream analysis. For the adult sample etrahypothalamic cells such as *PPP1R1B/DRD1/PDYN/DRD2/ADORA2A*+ cells (Nucleus accumbens) were removed from the anterior samples (preoptic and supraoptic) prior downstream analysis.

### Neuronal lineage analysis

Prenatal and adult hypothalamic neurons (CS22-Adult; n= 33,811) were integrated. *Seurat* was then used for dimension reduction using PCA and generation of a 3D UMAP. The scrattch.hicat package was used to generate 170 neuronal clusters from the adult hypothalamic neurons. Nuclei were identified based on expression of marker genes for each nuclei based on previously published studies (**table S3**) and/or colocalization in the Allen Brain Atlas (**fig. S4**).

*Monocle3*(*58*) was used to generate pseudotime lineages encompassing both prenatal and adult neurons. The Monocle3 trajectory generated 438 vertexes. Vertex 8 was chosen as the starting node as it contained the highest proportion of the earliest time point - the CS22 cells (66%). Using this pseudotime lineage, we identified branch points where the lineages diverge, and ‘leaves’ where the lineages terminate. Using the all_simple_paths function in Monocle3 and the *igraph* package, we were able to generate 2D lineage trees to represent the lineage. Gene expression was overlaid onto these lineage trees which showed the cells organizing into distinct lineages representing 10 anatomically distinct regions.

### Comparison across human/mouse and adult/prenatal neurons

Human adult hypothalamic neurons (n=34,662) and human fetal neurons (n= 16,451) were merged with mouse fetal samples (n=71,149) from Kim *et al*. 2020(*11*), and human adult data either from the whole mouse adult hypothalamus (n = 17,416) from Kim *et al*. 2020(*11*) and Chen *et al*. 2017(*5*), or specific nuclei from the adult mouse including Arcuate (n = 9,368) from Campbell *et al*. 2017(*4*), Preoptic nucleus (n=10,314) from Moffitt *et al*. 2018(*6*), Suprachiasmatic Nucleus (n=2,015) from Wen *et al*. 2020(*9*), Ventromedial Hypothalamus (n=12,062) from Kim *et al*. 2019(*7*), and the Lateral Hypothalamus (n=2,317) from Mickelsen *et al*. 2019(*8*). Louvain clustering was used to assign cells to 28 clusters. The distribution of cell types across these clusters was determined by first normalizing to the number of cells in each sample of the input.

The *POMC^+^* cluster was cleaned and then subclustered using louvain clustering. 11 clusters were determined based on distribution of arcuate marker genes (**table S7**). Transcription factors with high correlations to neuropeptides were determined for each sample type by co-occurance analysis using the *quanteda* package(*59*) in R and calculating the log likelihood significance for each pair of transcription factors and neuropeptides. From each of these matrices, the top transcription factors were determined as genes with co-expression above one standard deviation above the average of all correlations. If a gene appeared as a top transcription factor in all four matrices, they were considered conserved. Genes conserved in human were top transcription factors in both human datasets, but not in at least one of the mouse datasets. The transcription factors expressed in mouse, adult and fetal data were calculated accordingly.

### Comparison across brain regions during human development

Three fetuses from gestational week 18, 19 and 20 were chosen for their broad range of dissections covering the hypothalamus, three regions of the ganglionic eminence and five regions of the cortex. Adult neurons were filtered to include cells with at least 1000 reads (maximum 15000), number of genes 500 - 4000, a maximum 10 percent of reads mapping to mitochondrial genes and total UMI above 1000. The 95,107 cells passing these filters were integrated using the *Seurat* package in R with the same parameters as used for other datasets in this study. Cell type assignments were determined by a mixture of our work with the hypothalamus and cortex assignments previously reported(*57*). Three dimensional UMAP projections assisted in identification of multiple excitatory and inhibitory lineages as well as distinguishing ET and IT excitatory neurons and the three inhibitory neuron populations. Identification of IT and ET neuron types were further confirmed using *projectR* package and the *DeCoN* dataset(*60*). Seurat clustering at a resolution of 0.5 largely delineated major cell types. The Seurat function FindAllMarkers was used to both find top marker genes for each Seurat cluster as well as identifying brain region differences among Seurat clusters. Polar coordinate plots and Z-scores were calculated using the *volcano3D* (v2.0.1) package(*61*) and were used to identify top transcription factor differences among dividing progenitor cells. Genes selected for the heatmap in Figure 4E were a combination of top marker genes per Seurat cluster that were both similar across brain regions and as well different. Gene regulatory networks were reconstructed using the *Genie3* package(*32*). Networks were first built for each individual sample then edge weights were averaged across samples to identify region-specific networks. For summary analysis of networks in dividing cells, networks were combined by finding common edges and the resulting network was plotted using the *iGraph* package. Transcription factors with the greatest number of connections were displayed in white in figure 4F. Gene modules were created using K-means clustering starting with a K=100 and resulting in 28 groups, with gene eigenvalues calculated with the *WGCNA* package(*62*). Counts tables for both Genie3 and K-means clustering analysis were imputed using K nearest neighbors smoothing, where K = 5. We then compared gene regulatory networks across brain regions to identify region specific transcription factor drivers for each gene module. Transcription factors with poor correlation between the transcription factor expression and the gene module eigenvalue were discarded. Lists of transcription factor drivers per gene module word compared across brain regions and similarities and differences were reported.

**Fig. S1.**
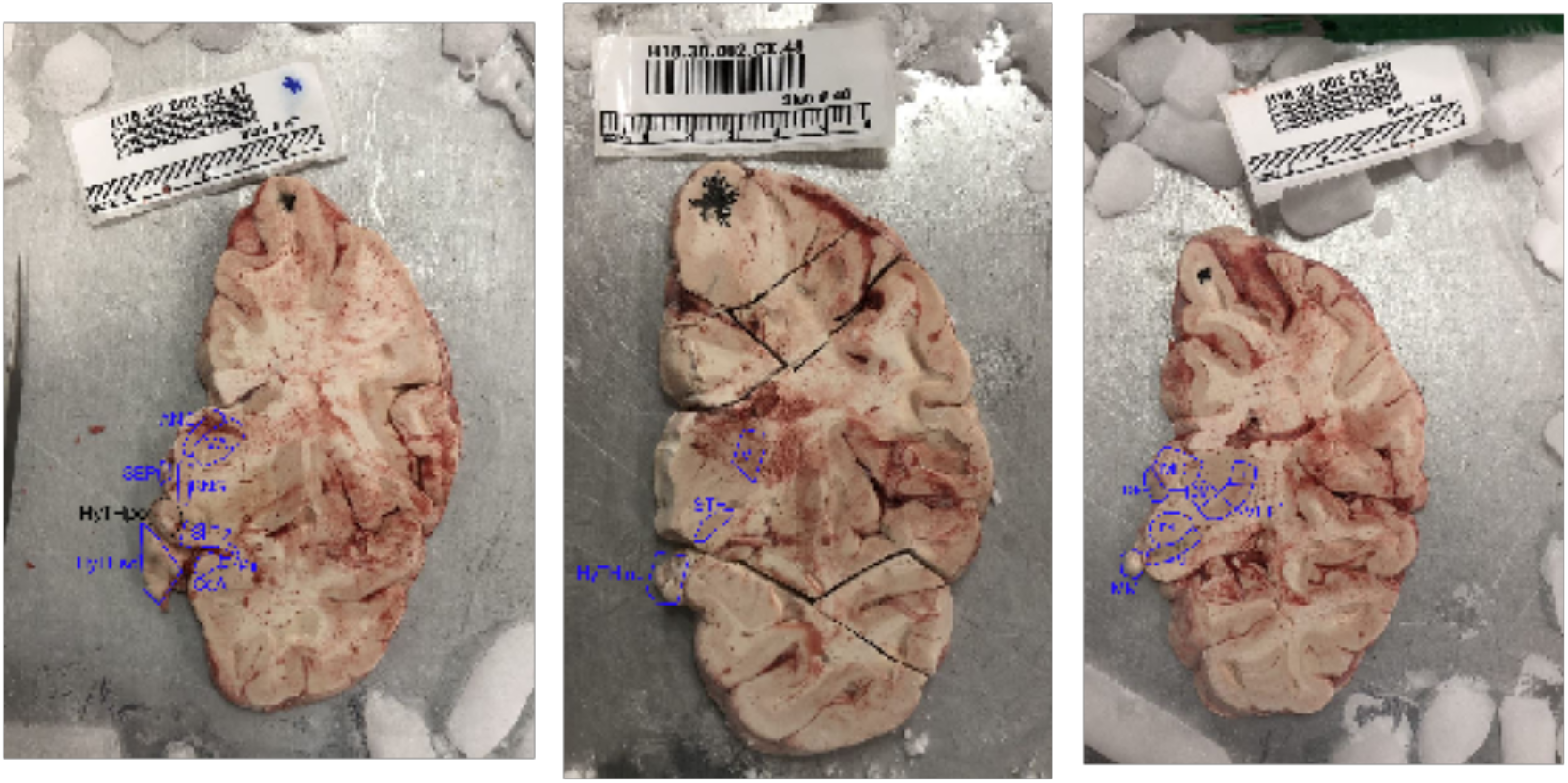
Dissection of the human adult hypothalamus. Coronal sections were taken from a 50 year old healthy male. Anatomical landmarks were used to locate the preoptic hypothalamus (HyTHpo), supraoptic hypothalamus (HyTHso), tuberal hypothalamus (HyTHtub) and mammillary hypothalamus (MM).

**Fig. S2.**
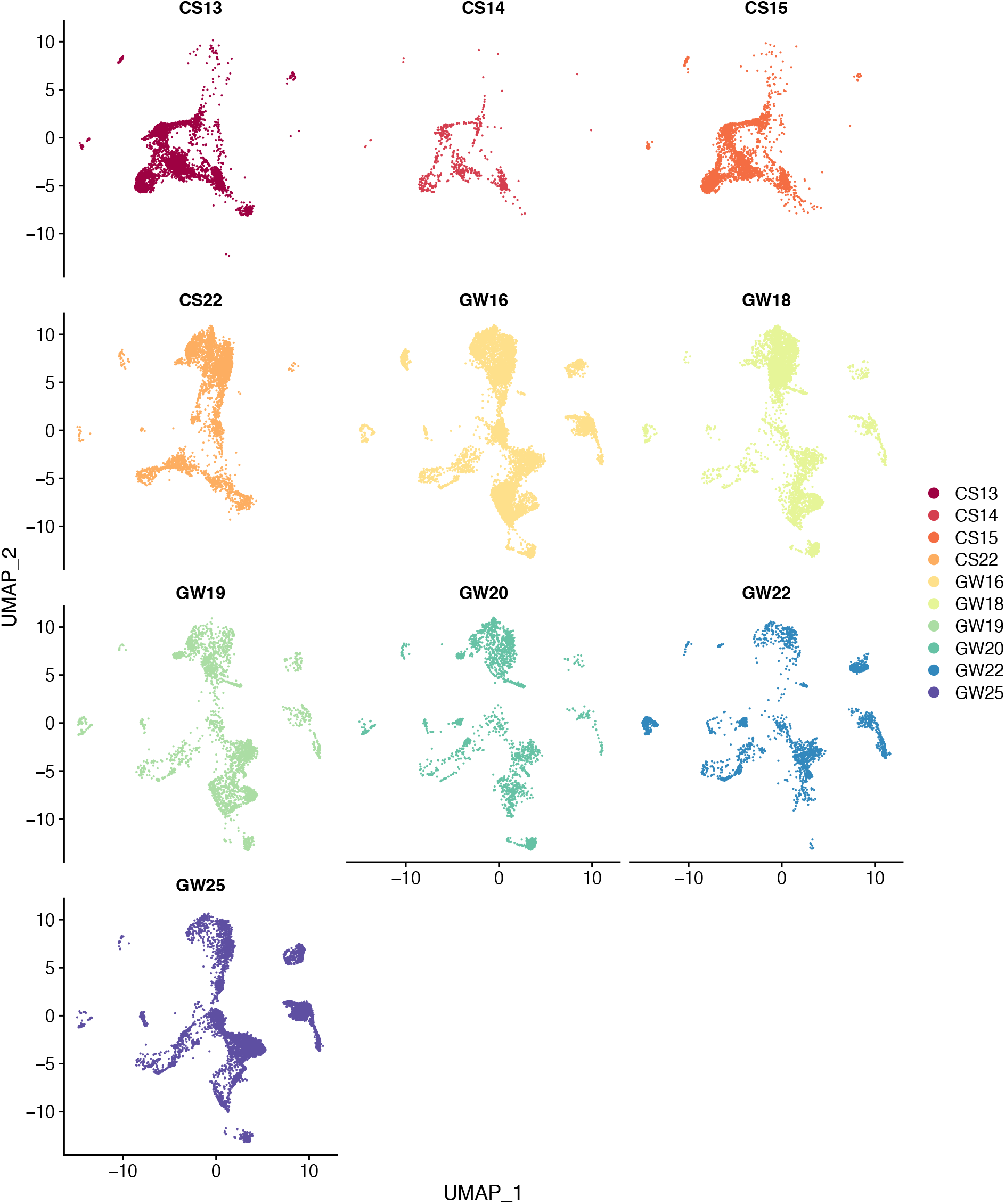
Distribution of samples throughout UMAP space. 2D UMAP representation of integrated human embryonic samples, each plot depicts the coordinates of cells in each sample.

**Fig. S3.**
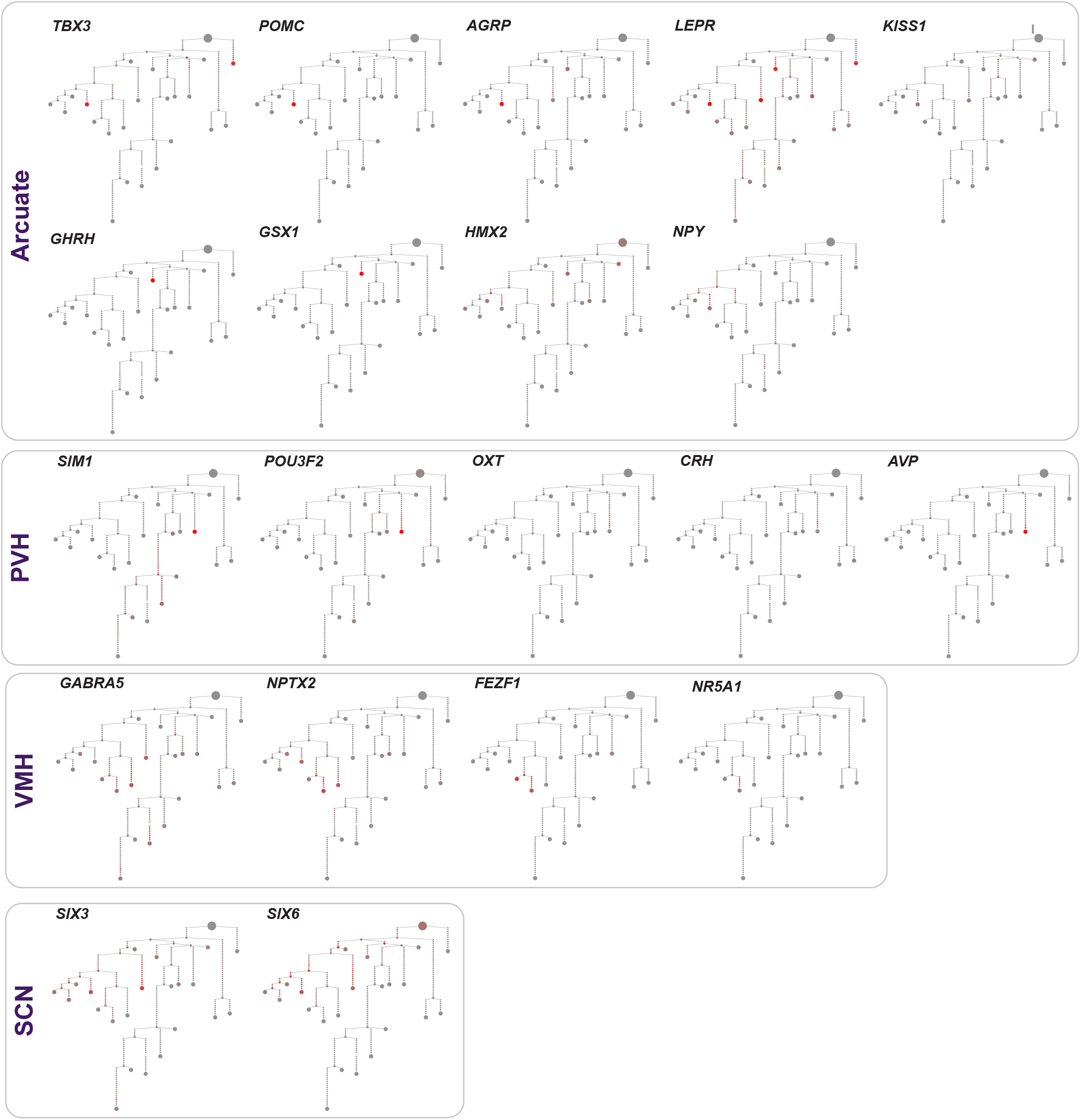
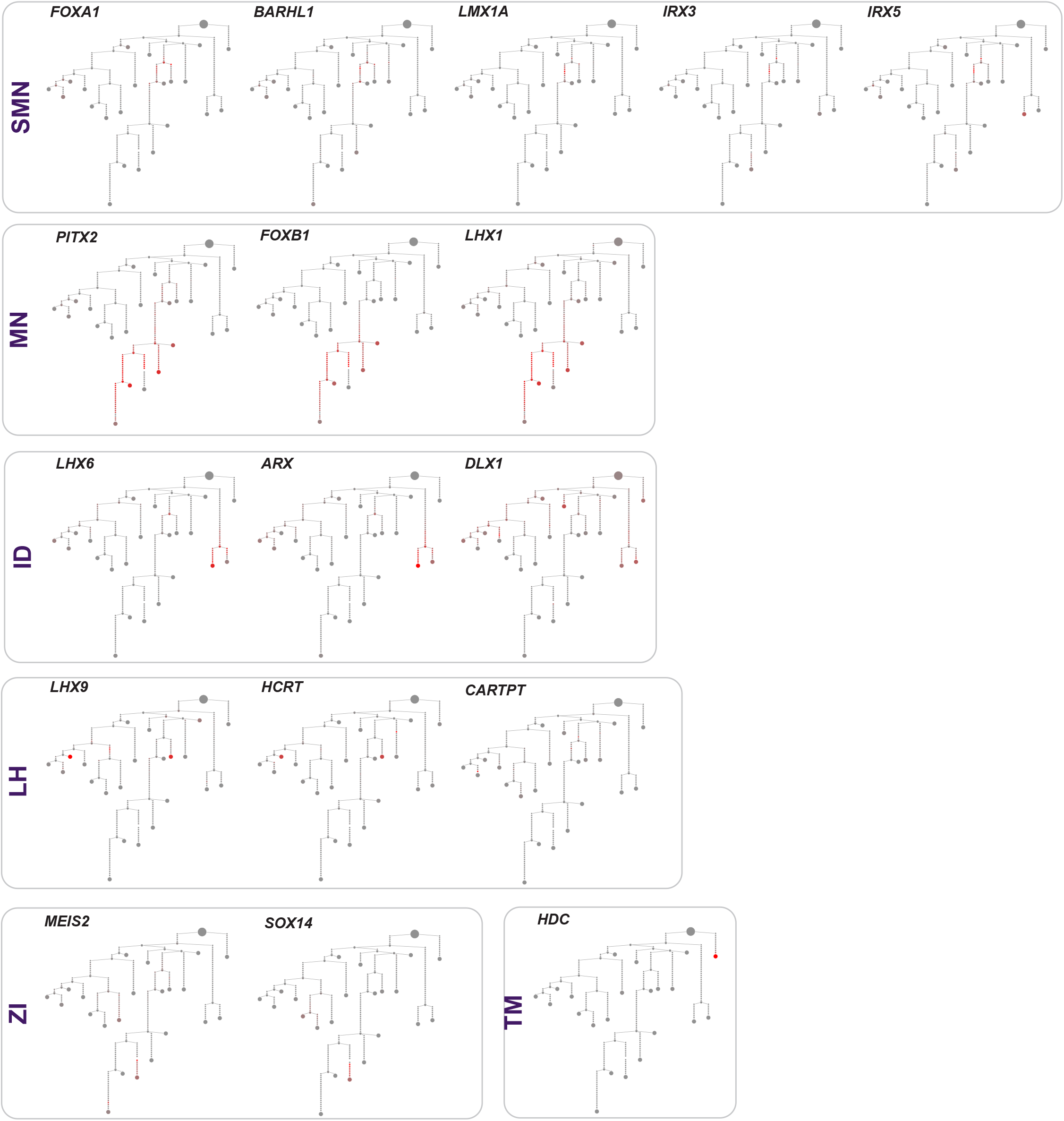
Expression of nuclei markers in embryonic and adult neuronal lineages. 2D lineage trees for all marker genes, with high expression indicated in red.

**Fig. S4.**
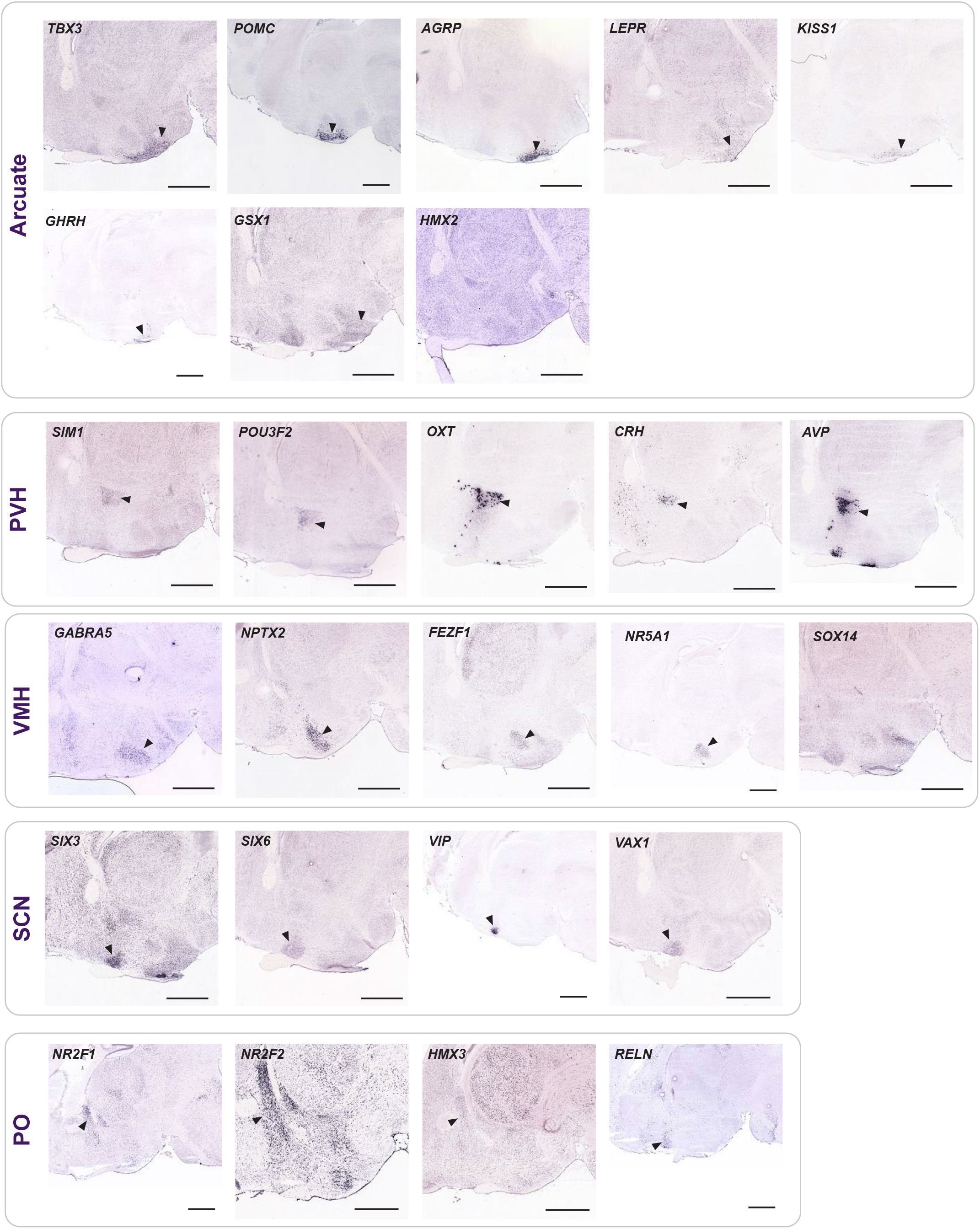
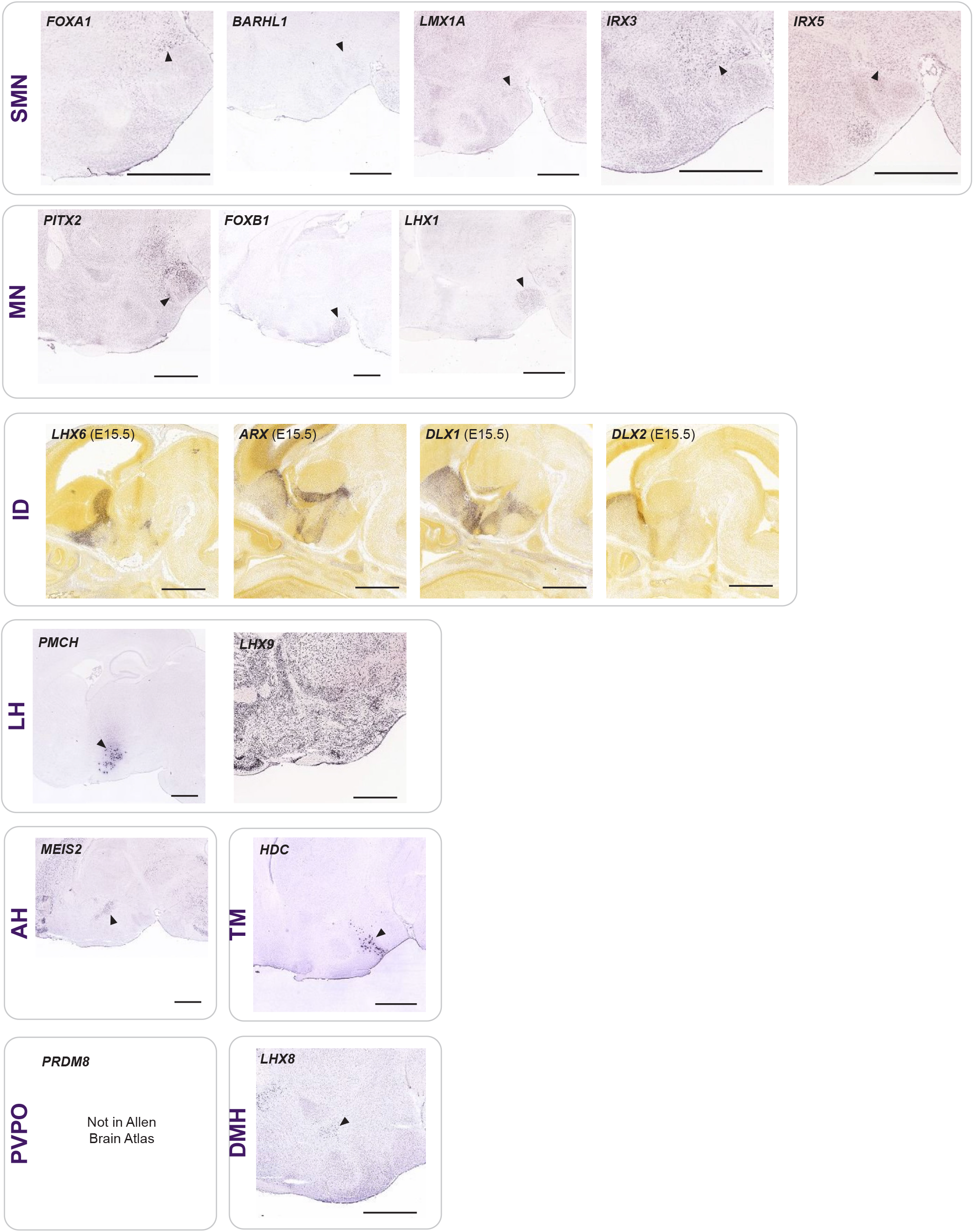
Localization of marker genes within the Allen Brain Atlas. In situ hybridization of marker genes used to discern neuronal lineages. Arrows show localization within the region of interest. Scale bar is 1000 microns.

**Fig. S5.**
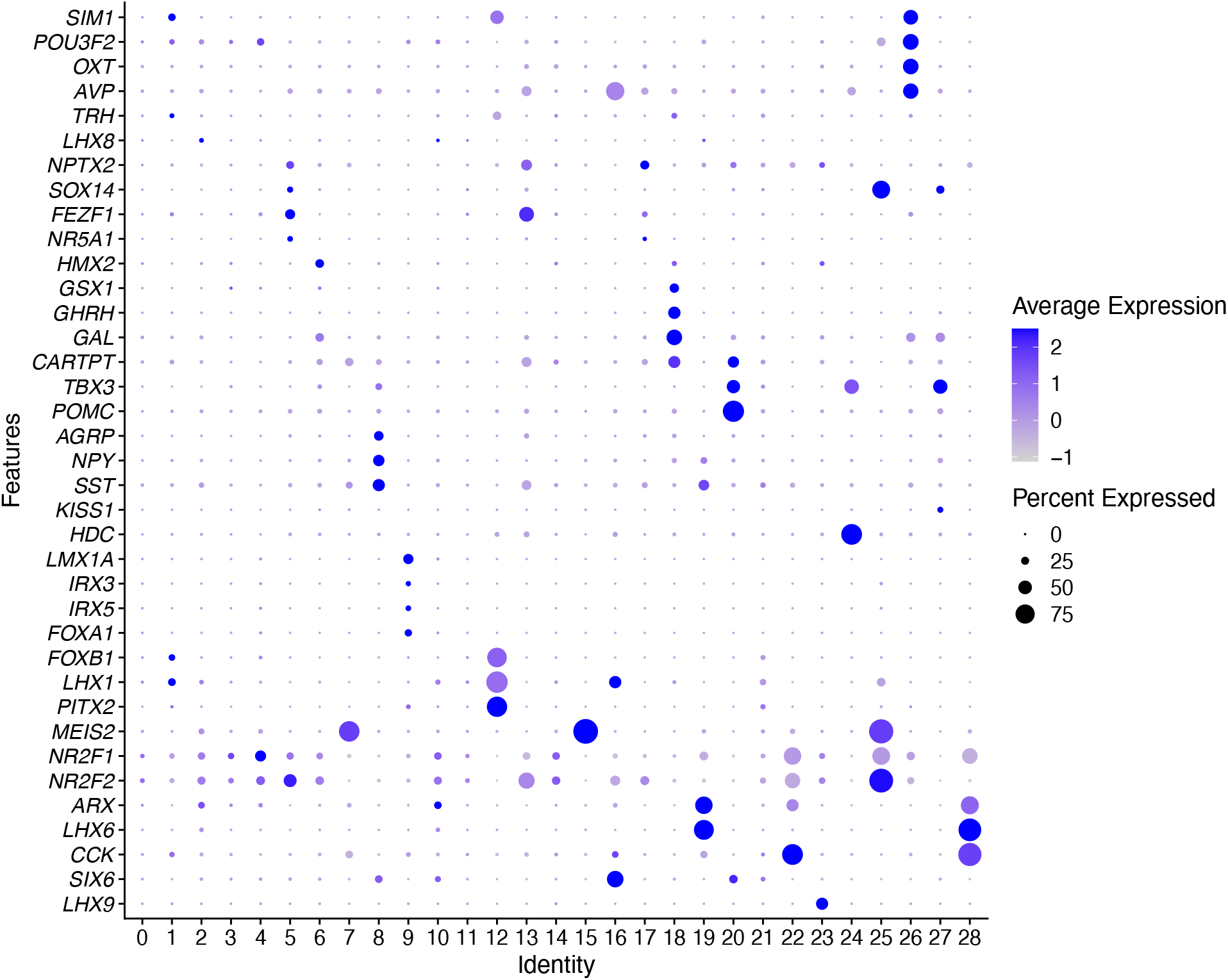
Expression of nuclei markers across species and development. Louvain clustering was performed on the integrated data from mouse and human, prenatal and adult samples. Nuclei specific markers are localized with Seurat clusters.

**Fig. S6.**
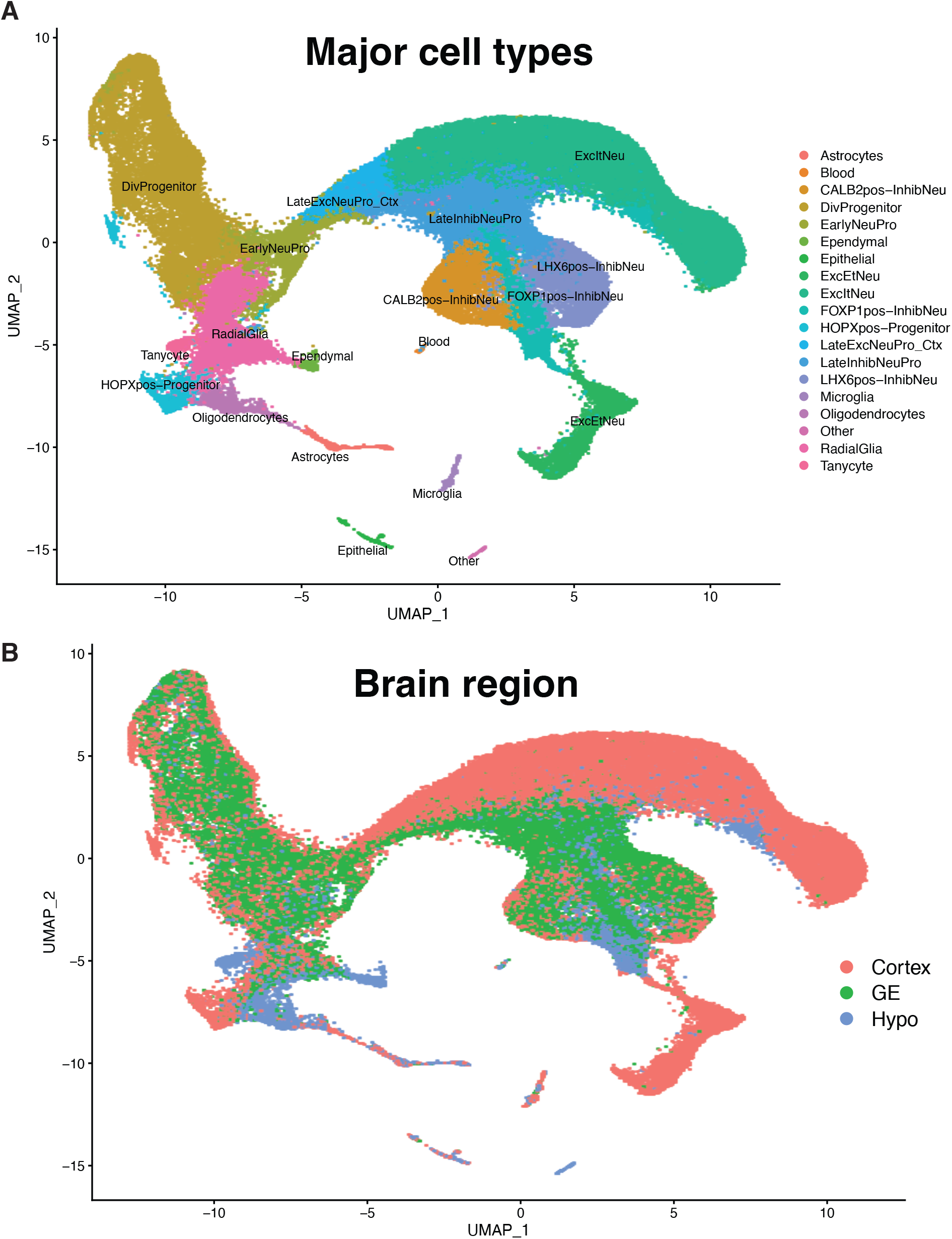
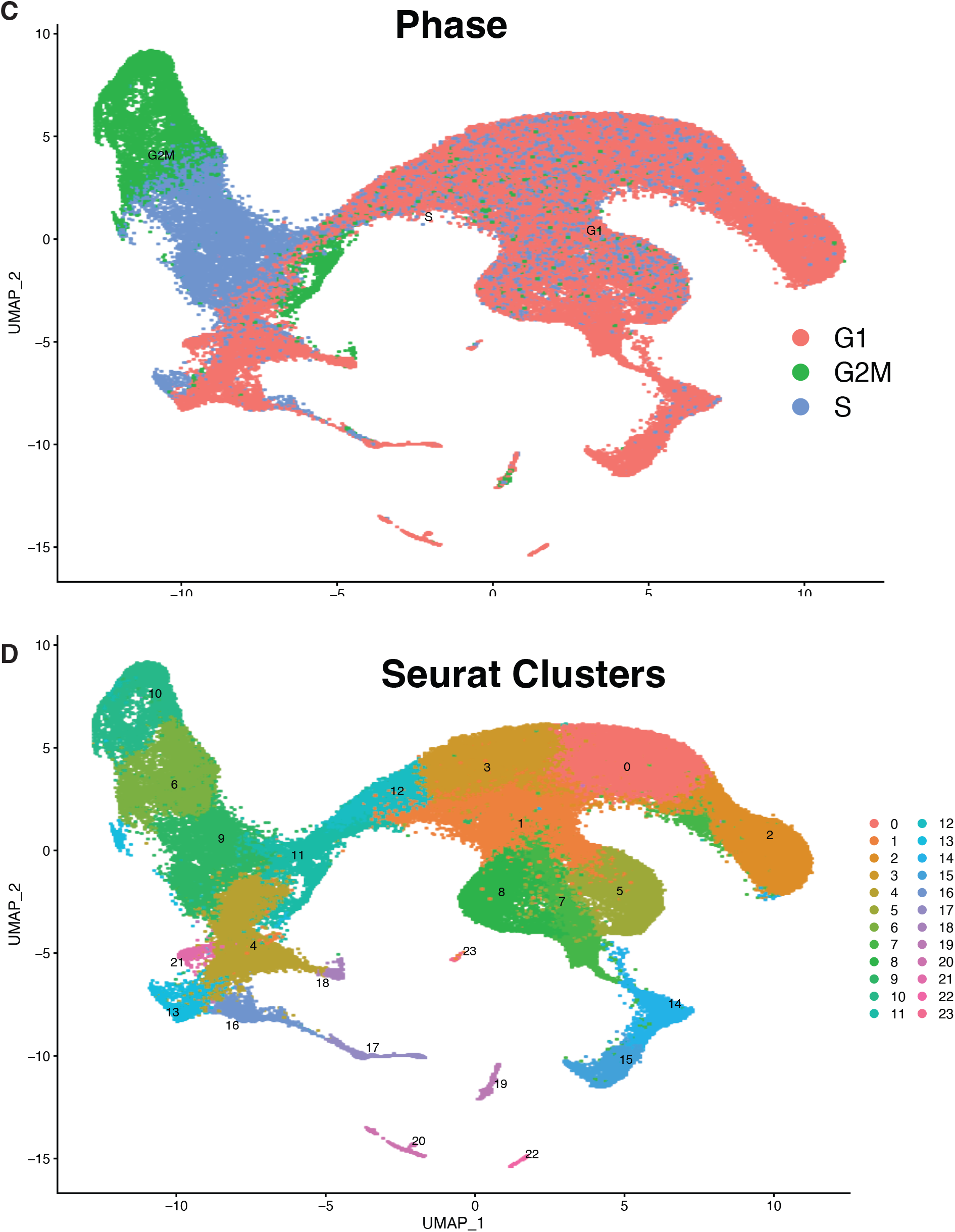
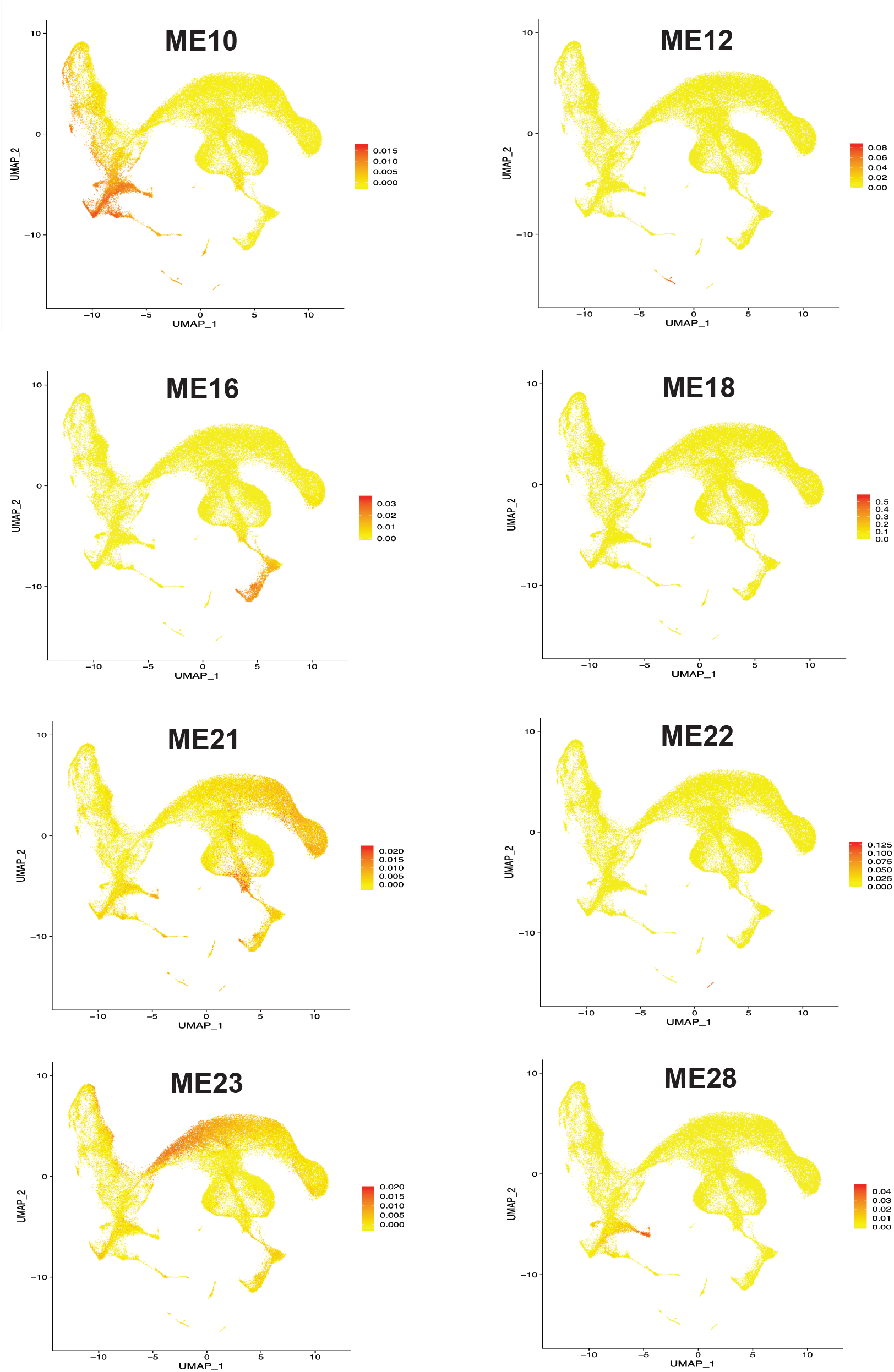
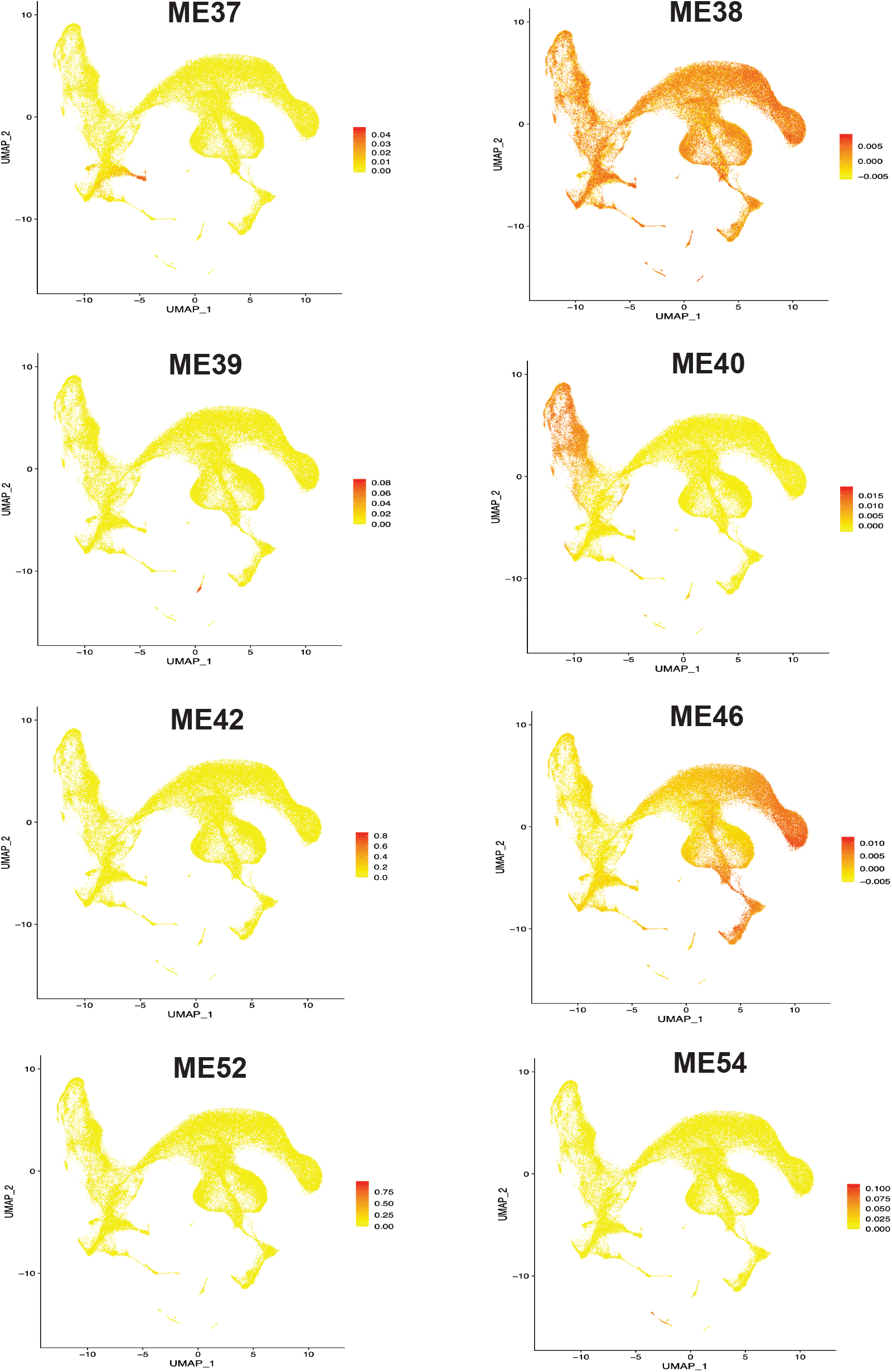
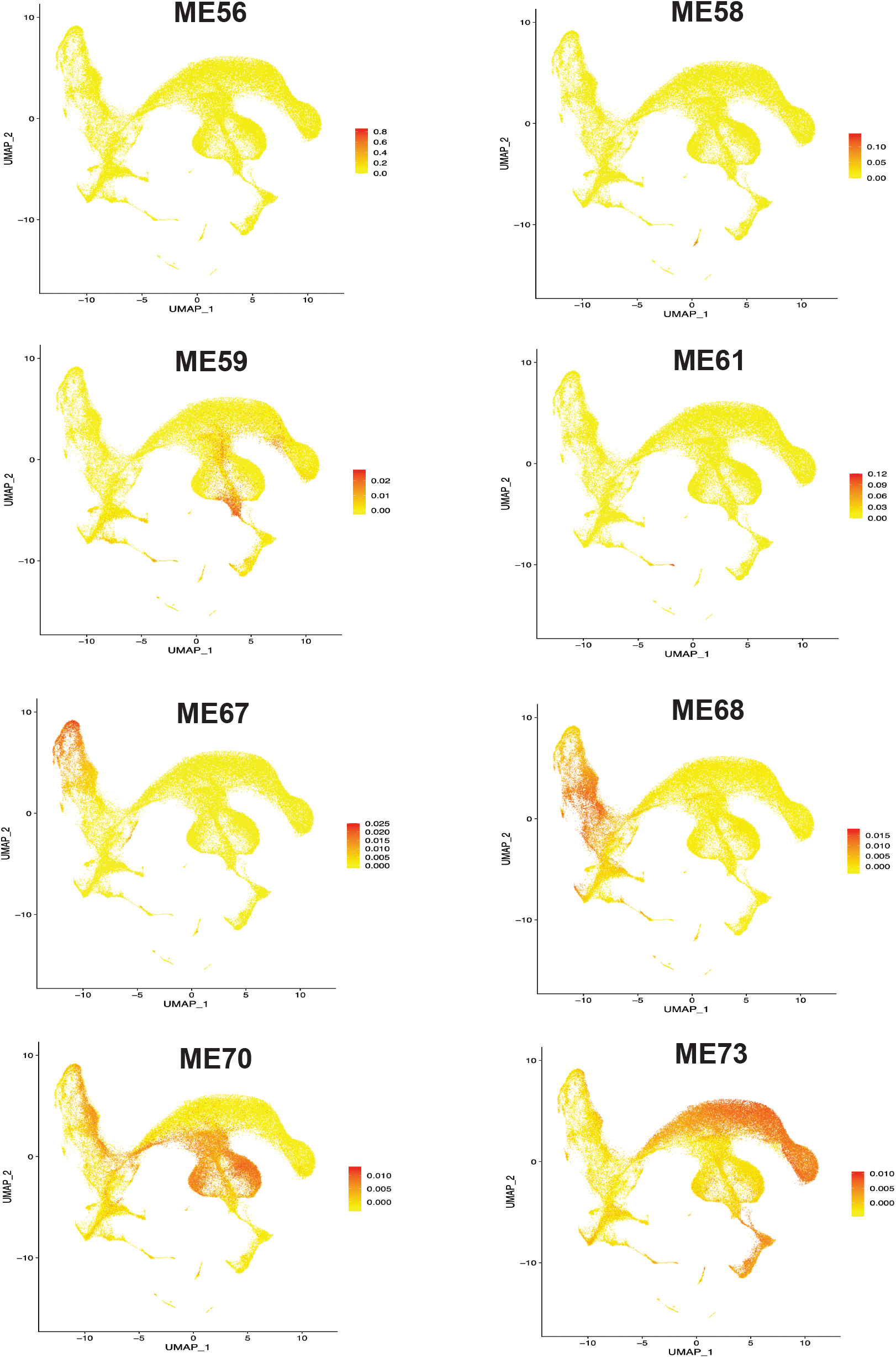
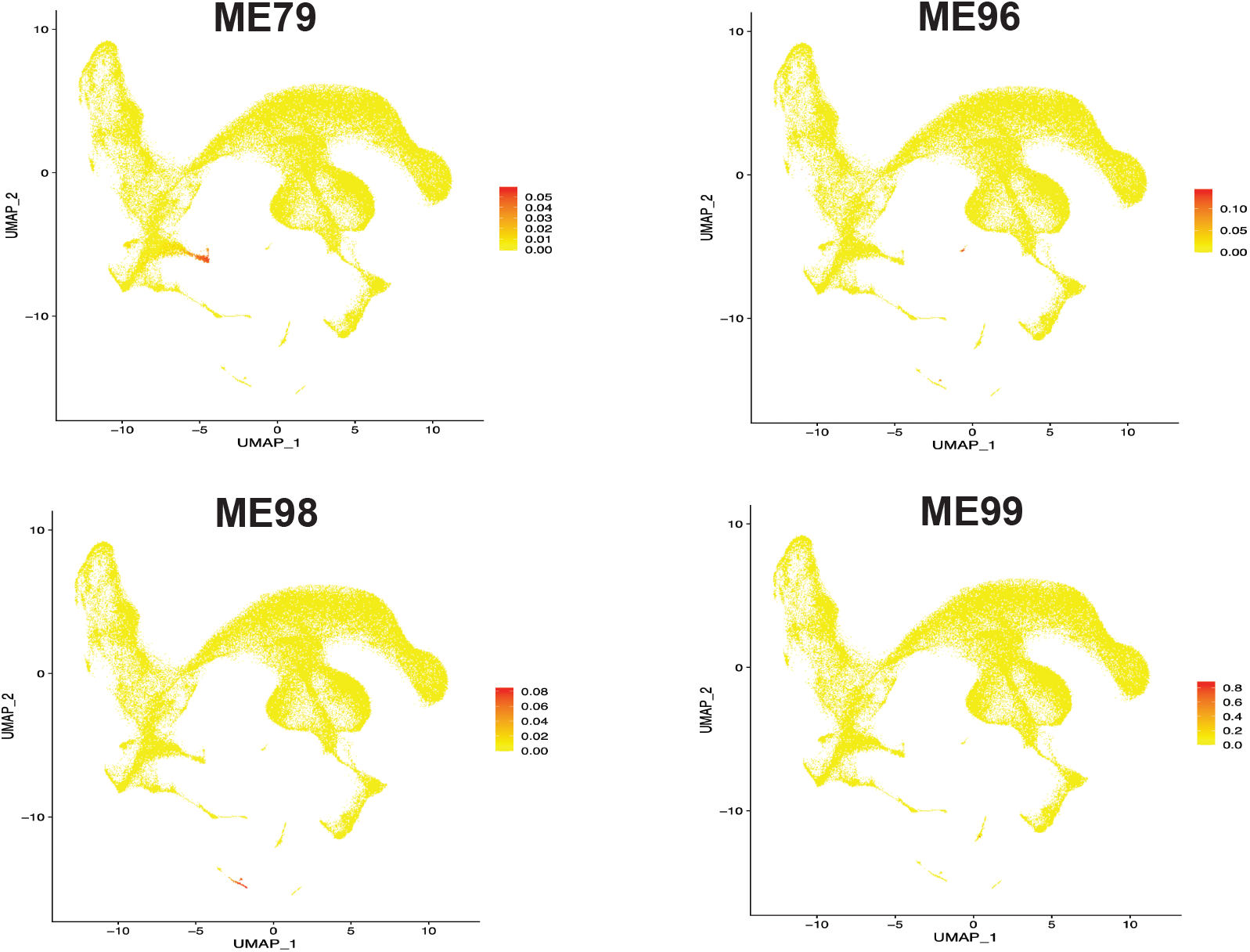
Gene module activity across cell-types and neuronal lineages. **A.** Major cell types determined by marker genes and boundaries defined by Seurat clustering. (3D UMAP) **B.** Integration of cells from across the hypothalamus, ganglionic eminence and cortex are presented in a 2 dimensional UMAP. **C.** Cell cycle stage as calculated using the Seurat program. (3D UMAP) **D.** Louvain clustering of cells calculated using the FindClusters function in Seurat, default resolution set to 0.5. (3D UMAP) **E.** Gene modules were created using imputed counts and grouping genes by K-means clustering. Eigengene values were calculated using the WGCNA package and these values were plotted for each gene module.

